# Anti-SARS-Cov-2 S-RBD IgG formed after BNT162b2 vaccination can bind C1q and activate complement

**DOI:** 10.1101/2022.04.24.489298

**Authors:** Anas H. A. Abu-Humaidan, Fatima M. Ahmad, Dima Awajan, Raba’a F. Jarrar, Nader Alaridah

**Affiliations:** Department of Pathology, Microbiology and Forensic Medicine, School of Medicine, The University of Jordan, Amman 11942, Jordan; Department of the Clinical Laboratory Sciences, School of Science, The University of Jordan, Amman 11942, Jordan; Department of Clinical Pharmacy and Therapeutics, Applied Science Private University, Amman 11931, Jordan

**Author notes:** Address correspondence to: Anas Abu-Humaidan M.D. Ph.D.

**Keywords:** COVID-19, vaccine, complement, C1q, C5b-9, BNTl62b2, BBIBP-CorV

## Abstract

**Introduction:** Activation of the classical complement pathway through C1q binding to immunoglobulins (Ig) contributes to pathogen neutralization, thus, the ability of Ig produced after vaccination to bind C1q could affect vaccine efficacy. In this study, we investigated C1q binding and subsequent complement activation by anti-spike (S) protein receptor-binding domain (RBD) specific antibodies produced following vaccination with either the mRNA vaccine BNT162b2 or the inactivated vaccine BBIBP-CorV.

**Methods:** Serum samples were collected in the period July 2021-March 2022. Participants’ demographic data, type of vaccine, date of vaccination, as well as adverse effects of the vaccine were recorded. The serum samples were incubated with S protein RBD-coated plates. Levels of human IgG, IgM, and C1q, that were bound to the plate, as well as formed C5b-9, were compared between different groups of participants.

**Results:** A total of 151 samples were collected from vaccinated (n=116) and non-vaccinated (n=35) participants. Participants who received either one or two doses of BNT162b2 formed higher levels of anti-RBD IgG than participants who received BBIBP-CorV. The anti-RBD IgG formed following either vaccine bound C1q, but significantly more C1q binding was observed in participants who received BNT162b2. Subsequently, C5b-9 formation was significantly higher in participants who received BNT162b2, while no significant difference in C5b-9 formation was found between the non-vaccinated and BBIBP-CorV groups. Formation of C5b-9 was strongly correlated to C1q binding, additionally, the ratio of formed C5b-9/ bound C1q was significantly higher in the BNT162b2 group.

**Conclusion:** Anti-RBD IgG formed following vaccination can bind C1q with subsequent complement activation, the degree of terminal complement pathway activation differed between vaccines, which could play a role in in the protection offered by COVID-19 vaccines. Further investigation into the correlation between vaccine protection and the ability of vaccine generated antibodies to activate complement is required.

## Introduction

COVID-19 caused by the severe acute respiratory syndrome coronavirus 2 (SARS-COV-2) was declared a pandemic in 2020 and has since infected more than 500 million people and resulted in over 6 million deaths. Vaccines produced against SARS-COV-2 had a major role in limiting the spread of infection and decreasing hospitalization[1]. There are currently over 150 vaccines in clinical development with around 10 vaccines approved for use, among them were the mRNA BNT162b2 developed by Pfizer–BioNTech and the inactivated vaccine BBIBP-CorV developed by Sinopharm. The two types of vaccines were found to offer various degrees of protection from COVID-19 in terms of mortality and hospitalization[2, 3], as well as varying humoral and T cell-mediated immune responses[4].

Vaccines initiate the formation of antibodies, prime T cell mediated immunity, and establish memory B- and T cell populations. The protection offered by antibodies can be ascribed to several effector mechanisms; one of which is binding to antigens on the surface of the pathogen to neutralize its ability to bind to host cells [5]. Several COVID-19 vaccines targeted the Spike (S) protein in SARS-COV-2, which is important for viral attachment to host angiotensin-converting enzyme 2 (ACE2) and entry into host cells [6–8]. Some of the antibodies produced following vaccination against SARS-COV-2 bind the RBD of S protein, such antibodies are commonly referred to as neutralizing antibodies [9].

In addition to binding viral antigens though the Fab portion, the Fc portion of antibodies can interact with Fc receptors found on immune cells [10] and with C1q of the complement system [11]. The complement system is made of several proteins that are found in tissue and circulation; those proteins are activated in a cascade-like manner to yield fragments that propagate the immune response. Pattern recognition molecules such as C1q and Mannose-binding lectin (MBL) can opsonize pathogens and activate the classical and lectin pathways respectively. Subsequently, C4 and C2 are cleaved to form the C3 convertase, which cleaves C3 and leads to formation of the opsonin C3b as well as the anaphylatoxin C3a, which interacts with a variety of innate and adaptive immune cells. Finally, activation of the terminal pathway through C5 cleavage leads to the formation of another potent anaphylatoxin, C5a, and leads to C5b deposition on the pathogen surface, which initiates the formation of the pore forming complex C5b-9 [12].

In COVID-19, activation of complement can occur through the classical, lectin, and alternative pathways. Activation of complement is meant to be a protective immune response against SARS-COV-2, but aberrant activation of complement is thought to contribute to the deterioration of patients with severe disease, as evidenced by the elevated levels of complement activation fragments found in the sera and lungs of hospitalized patients [13].

While the role of complement in COVID-19 was investigated in several studies, less attention has been given to the role of complement in the effectiveness or efficacy of COVID-19 vaccines. As mentioned above, C1q can bind immune complexes (antibodies bound to their respective antigen) and activate the classical pathway of complement, but the level of C1q binding and complement activation can vary between antibodies, for example, a recent study indicated that the structural features, such as the glycosylation patterns of the Fc portion of IgG could affect FcɣR and C1q binding[14]. Therefore, this study aimed to assess C1q binding and subsequent complement activation by anti-RBD antibodies produced following vaccination with either the mRNA vaccine BNT162b2 or the inactivated vaccine BBIBP-CorV.

## Materials and methods

### Study design and population

This study was conducted in the period July 2021 to March 2022. The participants were recruited at the vaccination centers at the University of Jordan before receiving either their first, second, or booster vaccine doses. The vaccines included in the study were the inactivated vaccine BBIBP-CorV (Sinopharm) and the mRNA vaccine BNT162b2 (Pfizer-BioNTech), which were the two main vaccines provided in Jordan. The records for COVID-19 vaccination in Jordan were electronic, each participant received a text message of the date and type of vaccine they had, which allowed for accurate documentation of vaccine data. For each participant, the following data was recorded 1) demographics included age, sex, height, and weight. 2) Vaccination date, type, and doses. 3) Adverse events following vaccination.

The participants were mainly residents of the capital Amman, mostly Jordanian, and all were from the MENA region. The inclusion criterion was age ≥18 years of age. While exclusion criteria were 1) having used complement inhibition therapy within the last 3 months and 2) documented complement deficiency or other immunodeficiencies.

### Sample collection

Samples were collected in plain blood collection tubes with a gel separator, then immediately preserved at 4°C and allowed to clot for a maximum of 2 hours. Samples were then transported on ice for further processing. Serum was collected by centrifugation of clotted blood tubes for 15 minutes at 1500 × g and 4°C. Serum was directly aliquoted into sterile EP tubes and stored immediately at −80°C until analysis. Serum samples were thawed on ice on the day of the experiments. Repeat freeze–thaw cycles were avoided to prevent protein degradation.

### Relative quantification of anti-RBD immunoglobulins and complement proteins

The assay used for relative quantification of IgG, IgM, C1q, and C5b-9 employed an indirect enzyme-linked immunosorbent assay (ELISA), using 96-Well SARS-CoV-2 Spike protein RBD-Coated Plates (ACROBiosystems), which were already blocked with 2% Bovine Serum Albumin (BSA). The plates and serum from participants were brought to room temperature, then each well received 100 μl from a solution of 1% serum in phosphate-buffered saline with 0.1% Tween 20 (PBST) and incubated for 90 mins at 37° C. Some wells were incubated with PBST only and were used as controls. Plates were then washed three times with PBST and blocked with 120 μl of blocking buffer (2% BSA in PBST) for 30 min at 37°C.

For IgG and IgM measurement, plates were incubated with either horseradish peroxidase (HRP)-conjugated rabbit anti-human IgG polyclonal antibodies (Abcam), or HRP-conjugated rabbit anti-human IgM polyclonal antibodies (Novus) respectively, both at a concentration of 1:2000 in blocking buffer at 4° C overnight.

While for C1q and C5b-9 measurement, the wells were washed three times with PBST, then incubated with either rabbit anti-human C1q polyclonal antibodies (MyBioSource) or mouse anti-human C5b-9 monoclonal antibodies (Novus), respectively, at a concentration of 1:500 in blocking buffer at 4° C overnight. The next day, the wells were washed three times with PBST, then incubated with the following secondary antibodies for 3 hours at room temperature; HRP-conjugated goat anti-rabbit polyclonal antibodies (Novex Life Technologies) for C1q, and HRP-conjugated goat anti-mouse polyclonal antibodies (Abcam) for C5b-9.

After the incubation with HRP-conjugated antibodies, all plates were washed three times with PBST. The chemiluminescence signal in the plates was measured as previously described [15], briefly, 50 μl of the enhanced chemiluminescence (ECL) substrate (Promega) was added to each well for 5 minutes, after development of the signal, images of the plates were taken using Chemidoc (Bio-Rad). The image files were analyzed using Fiji/ImageJ [16]. The signal from each well was normalized to signal from the control wells which were incubated with PBST instead of 1% serum, which is presented in the figures as signal/noise (S/N) fold change.

### Ethical approval

The study protocol was approved by the Institutional Review Board (IRB) at UJ (Ref. No. 68/2021). In addition, the work was conducted according to the principles of Good Clinical Practice (GCP) that has its origin in the Declaration of Helsinki (64th World Medical Association General Assembly, Fortaleza, Brazil, October 2013). All collected data were treated with confidentiality. Participation in the study was voluntary. A written and signed informed consent was obtained from all participants who agreed to participate following a full explanation of the study objectives.

### Data Analysis

Data generated was organized in Microsoft Excel, and statistical analysis was carried out using GraphPad prism 8 software for analysis. Results were presented as (mean ± SD) unless stated otherwise. The Shapiro-Wilk test was first used to test the distribution of data, subsequently, the nonparametric Mann–Whitney U test was used for single pairwise comparisons between dose matched vaccine groups, while the nonparametric Kruskal–Wallis test followed by Dunn’s test was used for multiple pairwise comparisons between vaccine groups. The nonparametric Spearman correlation coefficient was used to denote the magnitude and direction of correlation between different variables. All statistical tests were two-tailed and a probability value (p) less than 0.05 was considered significant.

## Results

### Demographics and vaccine-related information

All participants were above 18 years of age, and none reported any health conditions. We divided the participants who did not receive a vaccine into two groups: those who either tested negative for COVID-19 using a PCR test or did not perform a PCR test (PCR(-)) (n=22), and those who tested positive for COVID-19 using a PCR test at any time before sample collection (PCR(+)) (n=13).

While participants who were vaccinated were divided by the type of vaccine and the number of doses they received into the following groups: one dose of the BNT162b2 vaccine (1DP) (n=29), two doses of the BNT162b2 vaccine (2DP) (n= 27), one dose of the BBIBP-CorV vaccine (1DS) (n= 21), two doses of the BBIBP-CorV vaccine (2DS) (n=24), and a third (booster) dose of BNT162b2 vaccine following two doses of either the BBIBP-CorV or BNT162b2 vaccine (3D) (n= 15).

**Table 1.** Participants’ demographics and adverse events following vaccination.

| **Variables<sup>a</sup> / groups** | **Unvaccinated**<br>**(n= 22)** | **Unvaccinated**<br>**previously**<br>**PCR+**<br>**(n= 13)** | **BNT162b2**<br>**one dose**<br>**(n=29)** | **BBIBP-**<br>**CorV one**<br>**dose**<br>**(n= 21)** | **BNT162b2**<br>**two doses**<br>**(n=27)** | **BBIBP-**<br>**CorV two**<br>**doses**<br>**(n= 24)** | **BNT162b2**<br>**booster dose**<br>**(n=15)** |
| --- | --- | --- | --- | --- | --- | --- | --- |
| **Age (years)** | 25.18 (19.96,<br>30.41) | 26.62 (18.67,<br>34.56) | 28.14 (23.34,<br>32.94) | 32.62 (26.55,<br>38.69) | 29.96 (25.29,<br>34.63) | 25.83 (23.01,<br>28.66) | 23.67 (21.49,<br>25.84) |
| **Gender (percent)** |  |  |  |  |  |  |  |
| **Males** | 45% | 8% | 52% | 38% | 63% | 62% | 20% |
| **Females** | 55% | 92% | 48% | 62% | 37% | 38% | 80% |
| **Height (cm)** | 167.4 (163.4,<br>171.3) | 164.7 (160.1,<br>169.2) | 169.7 (166.3,<br>173.1) | 167.2 (164.1,<br>170.4) | 172.7 (168.7,<br>176.6) | 173.3 (168.9,<br>177.6) | 166.8 (162.6,<br>171.0) |
| **Weight (kg)** | 67.80 (61.34,<br>74.26) | 66.31 (54.01,<br>78.60) | 69.85 (64.07,<br>75.64) | 67.00 (62.93,<br>71.07) | 74.31 (66.73,<br>81.88) | 72.25 (65.36,<br>79.14) | 59.27 (55.05,<br>63.48) |
| **Time since the**<br>**latest infection or**<br>**vaccine dose**<br>**(days)** | NA | 187.1 (133.3,<br>240.9) | 34.34 (23.66,<br>45.03) | 102.7 (82.87,<br>122.5) | 113.4 (80.46,<br>146.4) | 72.46 (60.47,<br>84.45) | 67.07 (29.83,<br>104.3) |
| **Local symptoms**<br>**after vaccination**<br>1. Pain<br>2. Swelling<br>3. Redness<br>4. Itch | NA | NA | 1. 73%<br>2. 19%<br>3. 8%<br>4. 15% | 1. 31%<br>2. 6%<br>3. 0%<br>4. 6% | 1. 90%<br>2. 0%<br>3. 0%<br>4. 10% | 1. 58%<br>2. 0%<br>3. 0%<br>4. 0% | 1. 87%<br>2. 0%<br>3. 7%<br>4. 0% |
| **Systemic**<br>**symptoms after**<br>**vaccination**<br>1. Fever<br>2. Fatigue<br>3. Headache<br>4. Chills<br>5. Vomiting<br>6. Diarrhea<br>7. Muscle pain<br>8. Joint pain | NA | NA | 1. 23%<br>2. 42%<br>3. 29%<br>4. 4%<br>5. 0%<br>6. 0%<br>7. 42%<br>8. 12% | 1. 13%<br>2. 31%<br>3. 13%<br>4. 6%<br>5. 0%<br>6. 0%<br>7. 13%<br>8. 13% | 1. 25%<br>2. 65%<br>3. 30%<br>4. 10%<br>5. 0%<br>6. 5%<br>7. 45%<br>8. 35% | 1. 17%<br>2. 8%<br>3. 17%<br>4. 8%<br>5. 0%<br>6. 0%<br>7. 8%<br>8. 8% | 1. 33%<br>2. 40%<br>3. 27%<br>4. 27%<br>5. 0%<br>6. 0%<br>7. 20%<br>8. 13% |

<sup>a</sup>Data is presented either as mean and (95% confidence intervals) or as percent of participants.

### Relative levels of anti-RBD IgG and IgM

Analysis of relative anti-RBD IgG levels in various groups of participants was done by comparing each group to the PCR(-) group as a control. Using Dunn’s multiple comparisons statistical test following Kruskal–Wallis test indicated that all groups had significantly higher anti-RBD IgG levels than the PCR(-) group (Figure 1a). To assess the difference in anti-RBD IgG formation between BNT162b2 and BBIBP-CorV vaccines, the relative level of anti-RBD IgG was compared in each vaccine group using the Mann-Whitney statistical test, the comparison revealed higher levels of anti-RBD IgG in the 1DP compared to the 1DS group (mean S/N fold change: 93.79± 22.17, vs. 41.85± 39.92, respectively, p<0.0001) as well as higher levels in the 2DP compared to the 2DS group (mean S/N fold change: 99.27± 24.84, vs. 54.91± 31.90, respectively, p<0.0001). There was no significant difference between one or two doses in either vaccine.

**Figure 1.**
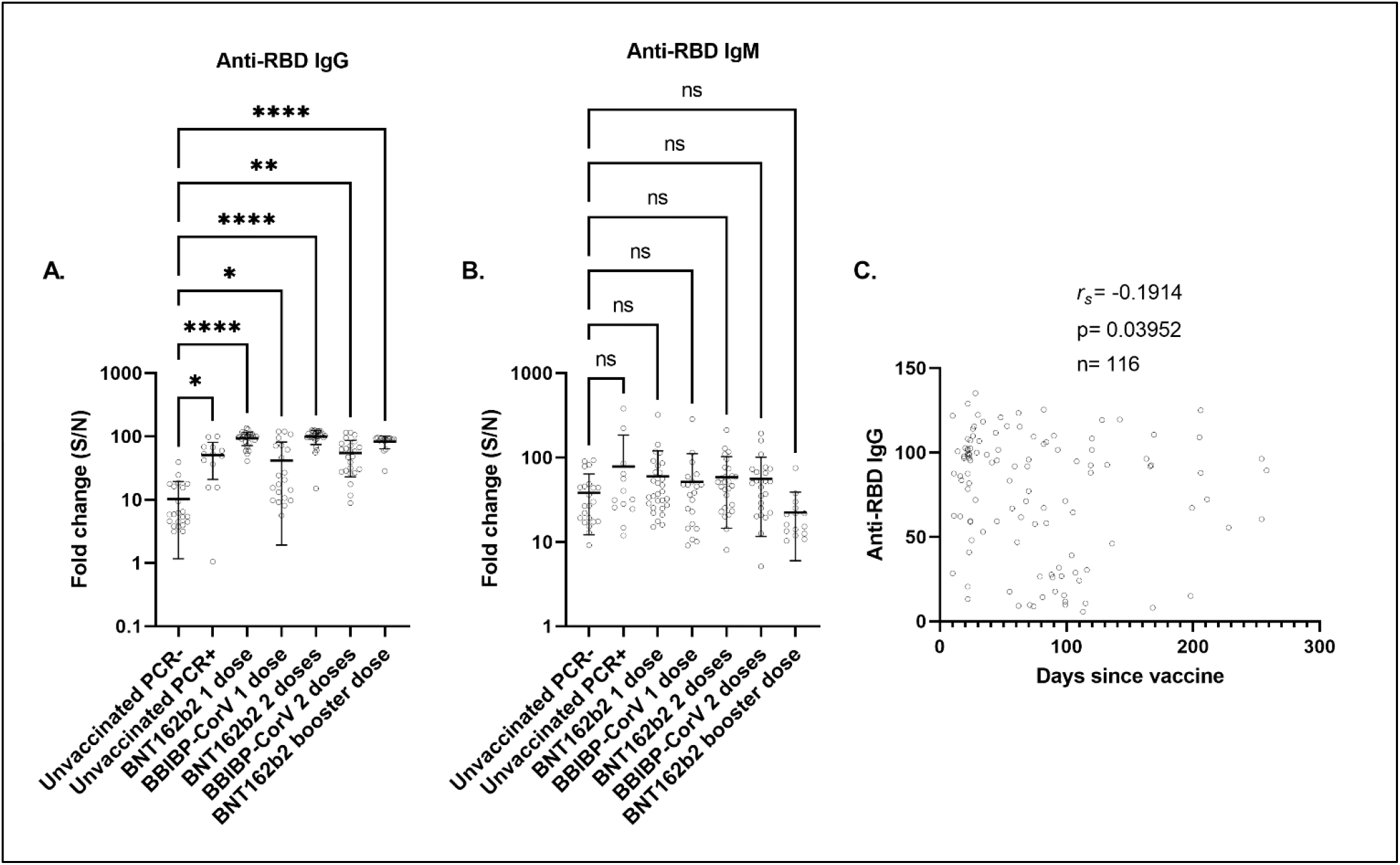
Measurement of anti-RBD IgG and IgM. Participants were divided by their vaccination status, type of vaccine, and dose of vaccine. Serum samples were incubated with RBD-coated plates; subsequently HRP tagged anti-human IgG or anti-human IgM antibodies were used to detect the presence of bound IgG or IgM, respectively. Fold change (S/N) represents the signal generated from wells incubated with serum divided by the signal in control wells incubated with PBST. (A) represents anti-RBD IgG levels while (B) represents anti-RBD IgM levels. (C) A scatter plot that correlates anti-RBD IgG levels to days since the vaccine in all vaccinated participants, spearman’s correlation coefficient (rs), p value, and number of participants are displayed on the plot. Each circle in the scatter plots represents one participant. Bold horizontal lines represent the mean of each group, while whiskers represent the standard deviation. Some error bars were clipped at the axis limit. ns P > 0.05, * P ≤ 0.05, ** P ≤0.01, **** P ≤ 0.0001.

The same statistical analysis was used to compare the groups of participants in terms of anti-RBD IgM levels, there was no significant difference in any of the groups when compared to the PCR(-) group (figure 1b).

Due to the setup of this study, the time between the latest vaccine dose and sample collection varied among participants, so we assessed the correlation between the time since the latest vaccine dose and the levels of anti-RBD IgG in vaccinated participants, we found a negative and weak yet statistically significant correlation using Spearman’s correlation coefficient (rs) (Figure 1c).

In aggregate, we confirmed that participants with a previous natural infection or vaccination with either dose of BNT162b2 or BBIBP-CorV had increased anti-RBD IgG. The anti-RBD IgG levels were significantly higher in the BNT162b2 vaccine groups. In addition, a significant difference in anti-RBD IgM was not found in any of the groups.

### C1q binding

Analysis of differences in C1q binding among groups of participants was done using Dunn’s multiple comparisons statistical test following Kruskal–Wallis test, and indicated that the 1DP, 2DP, 2DS, and 3D groups had significantly higher bound C1q than the PCR(-) group (Figure 2a).

**Figure 2.**
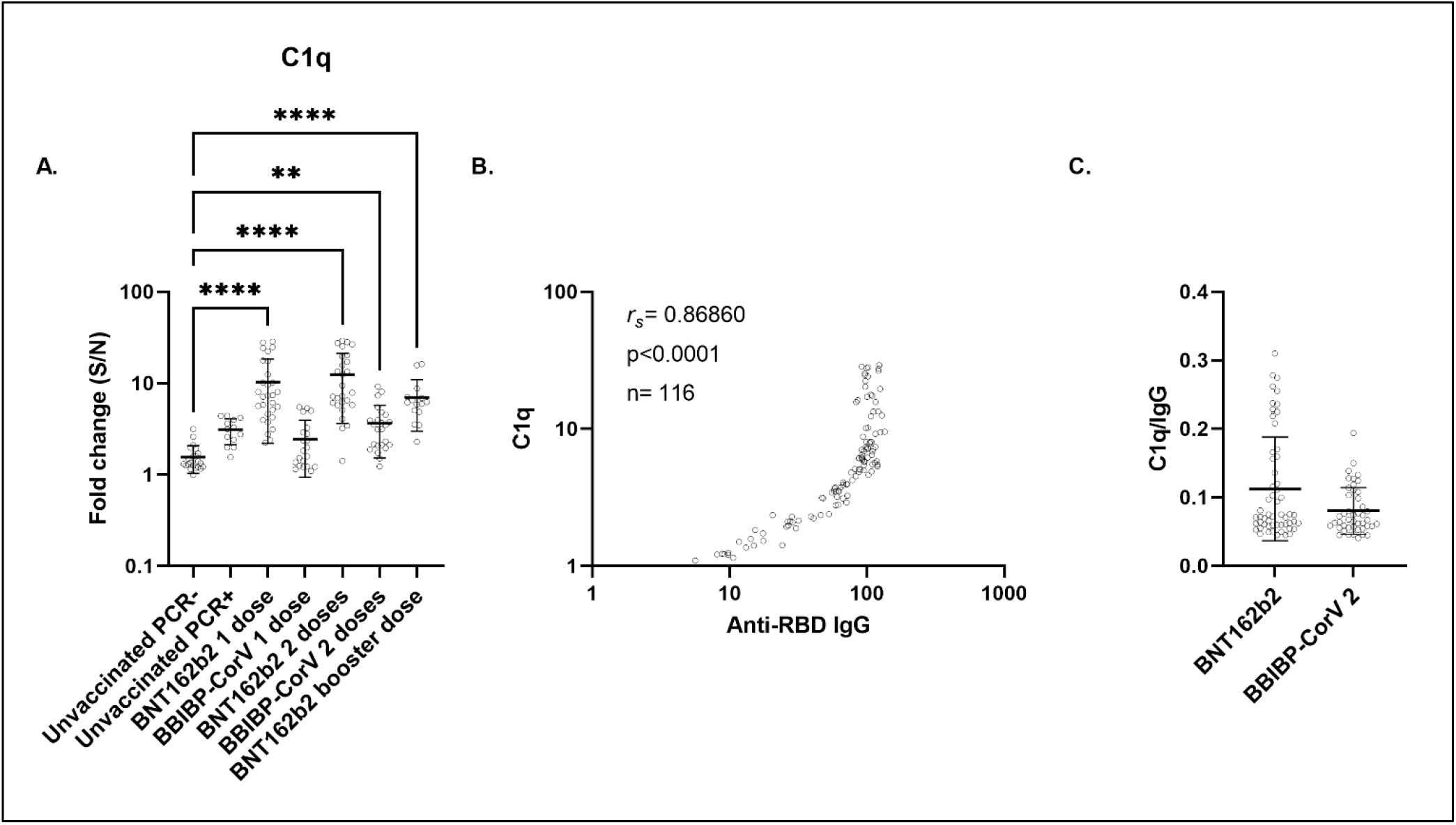
Measurement of C1q binding. Participants were divided by their vaccination status, type of vaccine, and dose of vaccine. Serum samples were incubated with RBD-coated plates; bound C1q was subsequently measured using an indirect immunoassay. Fold change (S/N) represents the signal generated from wells incubated with serum divided by the signal in control wells incubated with PBST. (A) represents bound C1q levels while (B) represents a log-log scatter plot that correlates C1q to anti-RBD IgG levels in all vaccinated participants, spearman’s correlation coefficient (*r*_s_), p value, and number of participants (n) are displayed on the plot. (C) Individual values of C1q per anti-RBD IgG in each participant were calculated and displayed in a scatter plot for the two types of vaccines. In scatter plots, each circle represents one participant, bold horizontal lines represent the mean of each group, while whiskers represent the standard deviation. Only significant pairwise comparisons are displayed, ** P ≤ 0.01, ****P ≤ 0.0001.

To assess the difference in C1q binding between BNT162b2 and BBIBP-CorV vaccines, the relative level of bound C1q was compared using the Mann-Whitney statistical test between each vaccine group, the comparison revealed higher levels of bound C1q in the 1DP compared to the 1DS group (mean S/N fold change: 10.27± 8.082, vs. 2.434± 1.496, respectively, p<0.0001), as well as higher levels in the 2DP compared to the 2DS group (mean S/N fold change: 12.39± 8.767, vs. 3.622± 2.101, respectively, p<0.0001). There was no significant difference between one or two doses in either vaccine.

To confirm that C1q was bound to the anti-RBD IgG, the correlation of bound C1q to IgG in all vaccinated participants (n=116) was assessed using Spearman’s correlation coefficient (rs). A strong and significant correlation was found between bound C1q and anti-RBD IgG (rs= 0.87, p<0.0001) (Figure 2b). Notably, the correlation between C1q and anti-RBD IgG was not linear after a certain point in the range tested (Figure 2b).

To address whether the increased binding of C1q in the BNT162b2 group was solely due to the increase in anti-RBD IgG level or due to increased ability to bind C1q, we examined the ratio of C1q/anti-RBD IgG in each participant in the BNT162b2 and BBIBP-CorV vaccine groups (Figure 2c). We found no significant difference in the ratio between the 2 groups suggesting that the amount of C1q bound per IgG did not differ significantly.

Taken together, the data indicate that the anti-RBD IgG formed following vaccination with either BNT162b2 or BBIBP-CorV were able to bind C1q. BNT162b2 led to the formation of more anti-RBD IgG and subsequently more C1q was bound.

### Activation of complement and formation of C5b-9

To assess whether binding of C1q led to activation of the terminal complement pathway, the formation of C5b-9 was used as an indicator. The analysis using Dunn’s multiple comparisons statistical test following Kruskal–Wallis test indicated that the 1DP, 2DP, and 3D groups formed significantly more C5b-9 than the PCR(-) group (Figure 3a). Notably, BNT162b2 vaccinated participants had a large range of terminal pathway activation, with a minimum, maximum, and mean fold change over control signal of 0.7593, 557.8, and 70.12 respectively in the 1DP group, and a minimum, maximum, and mean of 0.7727, 265.3 in the 2DP group.

**Figure 3.**
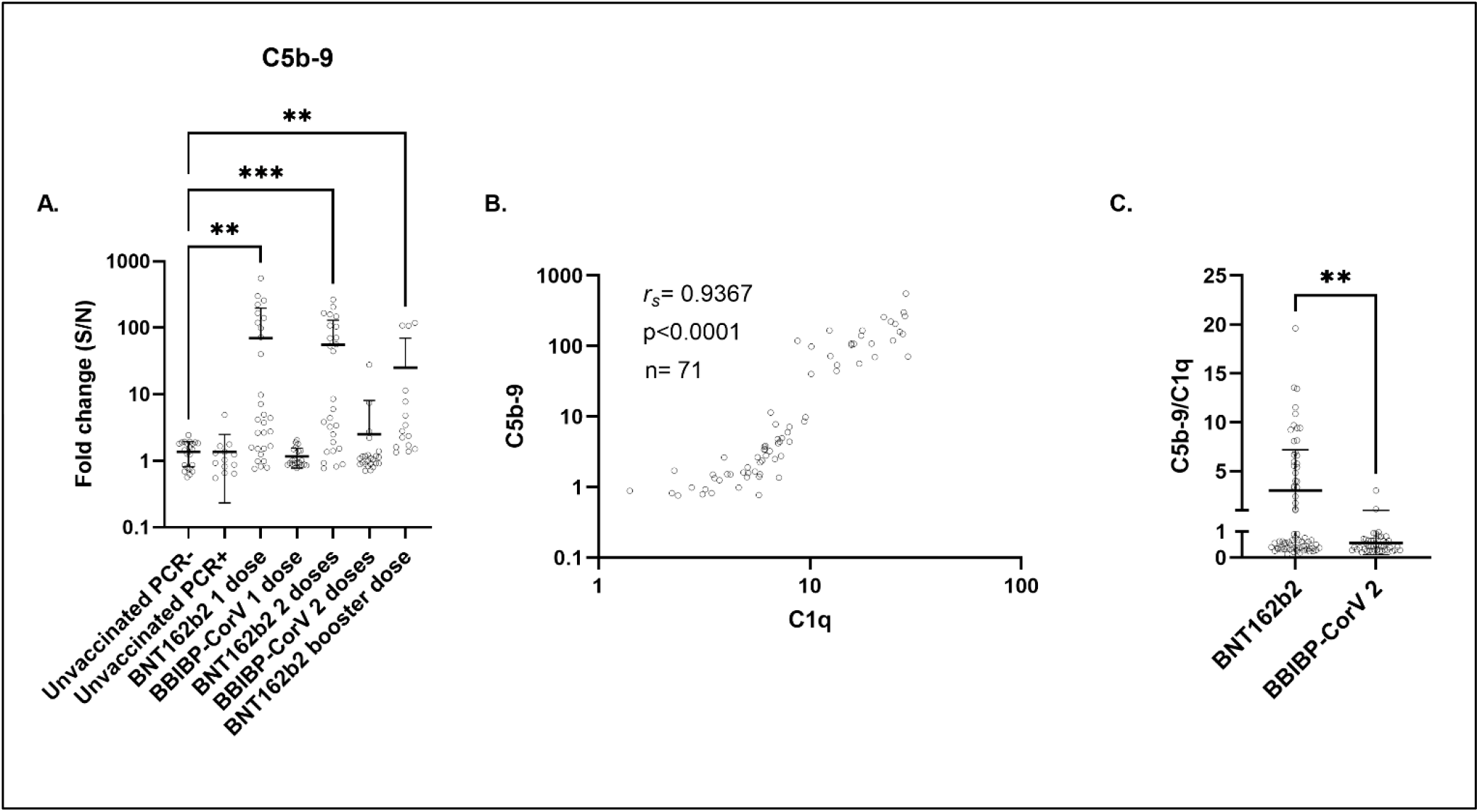
Measurement of C5b-9 formation. Participants were divided by their vaccination status, type of vaccine, and dose of vaccine. Serum samples were incubated with RBD-coated plates; the formation of C5b-9 was subsequently measured using an indirect immunoassay. Fold change represents the signal generated from wells incubated with serum divided by the signal in control wells incubated with PBST (S/N). (A) represents formed C5b-9 levels while (B) represents a log-log scatter plot that correlates C5b-9 to C1q levels in BNT162b2 vaccinated individuals, spearman’s correlation coefficient (rs), p value, and number of participants (n) are displayed on the plot. (C) Individual values of C5b-9 per C1q in each participant were calculated and displayed in a scatter plot for the two types of vaccines. In scatter plots, each circle represents one participant, bold horizontal lines represent the mean of each group, while whiskers represent the standard deviation. Some error bars were clipped at the axis limit. Only significant pairwise comparisons are displayed, ** P ≤ 0.01, *** P ≤ 0.001.

The correlation between C1q and C5b-9 formation was assessed in BNT162b2 vaccinated groups (1DP, 2DP, 3D) (n= 71), since they were the only groups that showed significantly higher C5b-9 than the PCR(-) group. A strong and significant correlation was found between bound C1q and C5b-9 (rs= 0.94, p<0.0001) (Figure 3b). Possibly indicating that the classical pathway activation following C1q binding is closely associated with activation of the terminal pathway and formation of C5b-9.

To address whether the increased C5b-9 formation in the BNT162b2 group was solely due to the increase in C1q binding or due to other factors, we examined the ratio of formed C5b-9/ bound C1q in each participant in the BNT162b2 and BBIBP-CorV vaccine groups (Figure 3c). We found significantly higher C5b-9 formation per C1q binding in the BNT162b2 group, this could be due to a more stable C1q binding leading to increased complement activation.

### Complement activation before and after BNT162b2 vaccination

To further confirm the findings of increased anti-RBD formation, increased binding of C1q, and increased complement activation following vaccination with the BNT162b2, we examined samples from 19 participants who provided a sample before and after the first dose of BNT162b2, the time between the dose and sample collection was (median 23 days, 95%CI 22-31). Additionally, 5 of the participants provided a second sample after the second dose, the time between the dose and sample collection was (median 19 days, 95%CI 14-37).

Vaccination with one dose confirmed the significant increase in anti-RBD IgG but not IgM (Figure 4. a, b), in addition, bound C1q was increased (Figure 4. c), and activation of the terminal pathway as evident by C5b-9 formation was increased (Figure 4. d).

**Figure 4.**
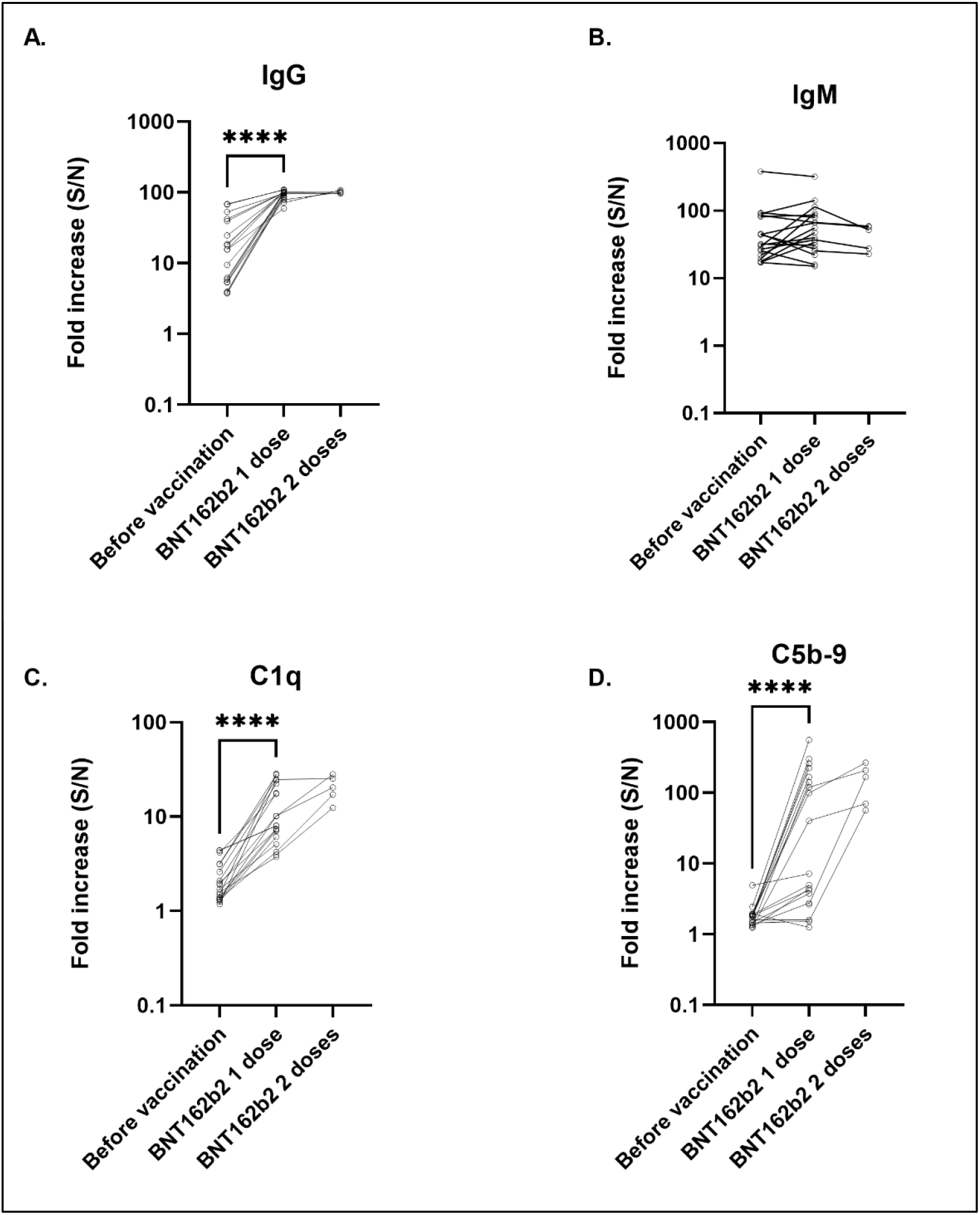
Complement activation before and after BNT162b2 vaccination. 19 participants provided samples before and after the first dose of BNT162b2, and 5 participants provided samples before and after the second dose BNT162b2. Serum samples were incubated with RBD-coated plates, bound IgG, IgM, C1q and formed C5b-9 were subsequently measured as described in methods. Fold change (S/N) represents the signal generated from wells incubated with serum divided by the signal in control wells incubated with PBST. The panels represent (a) bound IgG, (b) bound IgM, (c) bound C1q, and (d) formed C5b-9. Each circle in the scatter plots represents one sample, the circles connected by lines represent before and after samples from the same participant. Only significant pairwise comparisons are displayed, **** P ≤ 0.0001.

## Discussion

Vaccination confers immunity through several mechanisms one of which is antibody production. But other functional abilities of antibodies contribute to conferring protection against infection as well, such as activation of complement through C1q binding. This study aimed to assess the ability of anti-RBD antibodies formed after vaccination with either BBIBP-CorV or BNT162b2 in fixing C1q and activating the complement system.

The assay we employed for relative quantification of antibodies and complement proteins used signal from PBST treated wells as the background signal, and signal from wells treated with 1% serum as the true signal, the choice of 1% serum was guided by current literature and was confirmed through optimization assays. Although we cannot rule out unspecific binding of IgG and IgM especially since the signal in the unvaccinated PCR(-) group was significantly higher than the background, the group of unvaccinated PCR(-) participants served as the control group for all statistical analysis purposes in this study, effectively discounting the unspecific binding if present.

The results indicate significantly higher anti-RBD IgG in previously infected participants (PCR(+))when compared to the PCR(-) group, the median time since the last positive PCR test in the PCR(+) group was 182 days, indicating that anti-RBD IgG can persist for over 6 months, in accordance with a recent report where those antibodies persisted for around 20 months following a positive PCR test [17]. In this study, samples were collected from participants just before getting either the first, second, or booster doses, this led to variability in the time of sample collection after vaccination, either due to participants not adhering to the proposed dosing schedule or getting infected in between doses causing an extension of the period until the next dose. Nevertheless, in the time period in which we collected the samples, we found that the time of sample collection after vaccination had a minimal effect on anti-RBD IgG levels, as indicated by the weak negative correlation between the two, additionally, the comparison between the BBIBP-CorV or BNT162b2 clearly showed that the 2DP group had almost twice as much anti-RBD IgG than 2DS, although the time since the latest vaccine dose in the 2DP and 2DS groups was 113.4 days and 72.46 days, respectively.

Examination of anti-RBD IgM revealed no significant differences between any of the groups tested, which is probably due to the time point at which samples were collected. Anti-RBD IgM levels are significantly decreased at 6 months post infection[18], in addition, vaccination does not elicit formation of anti-RBD IgM as natural infection [19], both reasons account for our finding of no significant difference in anti-RBD IgM levels. This study also indicated that anti-RBD IgG but not IgM was more important in complement activation following vaccination, where C5b-9 strongly and significantly correlated to the amount of bound C1q, which in turn strongly correlated to the amount of bound anti-RBD-IgG but not IgM.

Binding of the Fc portion of IgG to C1q and subsequent complement activation has been shown to increase the neutralizing ability of antibodies in several types of infections[20–22]. The findings of this study indicated that anti-RBD IgG formed following vaccination with BNT162b2 and BBIBP-CorV bound C1q, as shown by the strong and significant correlation between the two in both vaccine groups. In addition to activating complement, binding of C1q to antibodies can improve virus neutralization in other less recognized mechanisms, for example, C1q binding to antibodies against West Nile virus was shown to reduce the stoichiometric requirements for neutralization[21], indicating that C1q binding on its own, even without subsequent complement activation can enhance neutralization.

We believe that the neutralizing ability of antibodies formed against SARS-COV-2 should be investigated in the presence and absence of C1q and other complement components, especially in light of advancements that allow the manipulation of the Fc portion of monoclonal antibodies to enhance effector functions[23], a process that is relevant in vaccine production as well[24].

When examining C1q binding in different vaccine groups, data indicated that serum from participants who took one or two doses of BNT162b2 bound more C1q than serum from participants who took one or two doses of BBIBP-CorV. While the ratio C1q/anti-RBD IgG did not differ significantly between the two vaccine types, the ratio C5b-9/C1q did. This finding could be explained by the difference in C1q binding stability in the presence of IgG oligomers, indeed, several studies reported on the enhanced ability of IgG oligomers, especially hexamers, in binding C1q and initiating complement activation[25, 26]. Notably, the formation of IgG oligomers and subsequent complement binding and activation depends to some extent on the nature of the IgG, in addition to antigen distribution on the surface[25, 27].

It was recently shown that anti-RBD IgG from SARS-COV-2 infected individuals can bind C1q and activate complement, and the amount of complement activation was associated with disease severity[28]. In our study, bound C1q and formed C5b-9 were both higher in the PCR(+) compared to the PCR(-) group but did not reach statistical significance, probably since participants were not hospitalized and did not have an active infection at the time of sample collection. Another recent study investigated Fc glycosylation patterns in IgG formed following BNT162b2 vaccination or natural infection, and found a unique proinflammatory Fc composition following BNT162b2 vaccination[14]. Such data emphasizes the importance of identifying unique effector functions in antibodies formed following vaccination, and the differences those effector functions have in the protection offered by various vaccines.

In conclusion, this study demonstrated increased C1q binding to anti-RBD IgG with subsequent complement activation in individuals receiving one, two, or a booster dose of BNT162b2, compared to individuals who receive one or two doses of BBIBP-CorV. Further studies are required to elucidate the relationship between the neutralizing ability of antibodies formed following COVID-19 vaccination and their ability to bind C1q and activate complement.

## Author contributions

AHAA contributed to study conception, study design, acquisition, analysis and interpretation of data, manuscript writing, and revision.

FA contributed to study design, acquisition, analysis, and interpretation of data, manuscript writing, and revision.

DA contributed to acquisition and interpretation of data, manuscript writing, and revision.

RJ contributed to acquisition and interpretation of data, manuscript writing, and revision.

NA contributed to acquisition and interpretation of data, manuscript writing, and revision.

## Acknowledgments

We are grateful for the help provided by the staff at UJ vaccine centres, especially Mr. Arafat Sawalha for facilitating sample collection.

